# An exceptionally preserved euarthropod with unique feather-like appendages from the Chengjiang biota

**DOI:** 10.1101/2021.01.22.427827

**Authors:** Dayou Zhai, Mark Williams, David J. Siveter, Derek J. Siveter, Thomas H.P. Harvey, Robert S. Sansom, Huijuan Mai, Runqing Zhou, Xianguang Hou

## Abstract

Micro-CT scanning of the Cambrian euarthropod *Chuandianella ovata* reveals unprecedented three-dimensional soft-part details. It has an elongate uniramous antennule and a short uniramous second appendage, followed by ten homonomous biramous appendages, each comprising a short paddle-shaped exopod and a unique feather-like endopod with at least 27 podomeres each of which bears a long blade-like endite with a short terminal seta. Based on its carapace and previously known soft-part anatomy *C. ovata* was compared with the Burgess Shale mandibulate euarthropod *Waptia*. However, *Waptia* has recently been shown to bear specialized head appendages that are interpreted as a mandible and maxillula, posterior to which are four appendages each with five-segmented endopods. In contrast, we interpret *Chuandianella* as an ‘upper’ stem-group euarthropod that possessed neither a differentiated mandible nor a maxillula. *Chuandianella* further demonstrates that early Cambrian ‘upper’ stem-group euarthropods were experimenting with a range of different limb arrangements and morphologies.

## Introduction

During the Cambrian Period bivalved arthropods formed a numerically abundant and widespread component of marine ecosystems in benthic, nektobenthic and pelagic settings (*Williams et al., 2007, 2015*). They are among the most abundant animals in the Chengjiang Lagerstätte of China (*Zhao et al., 2012; Hou et al., 2017*) and the North American Burgess Shale Lagerstätte (*Briggs et al., 1994*). Many of these arthropods are species that have been assigned to Bradoriida Raymond, 1935, a group common worldwide and with very rare exceptions known only from their bivalved carapaces (*Williams et al., 2007*). Where they preserve soft-part anatomy in many cases conventional classifications based on carapace morphology break down. The markedly different arthropod body plans revealed beneath the bivalved carapace include stem euarthropods (*Zhai et al., 2019a*), and mandibulates (*Vannier et al., 2018*) including stem pan-crustaceans (*Zhai et al., 2019b*).

Recent Micro-CT scanning of fossil arthropods has in many cases revolutionised our understanding of their soft-part anatomy. This has become particularly apparent in the study of the Cambrian (Series 2, Stage 3) Chengjiang biota arthropods of China (e.g., *Liu et al. 2020*), which preserve components of their original three-dimensional soft-part anatomy, including fine details of appendages (e.g., *Zhai et al., 2019a,b; Liu et al., 2020*). Here we describe newly micro-CT analysed specimens of the Chengjiang bivalved arthropod *Chuandianella ovata* (*Li, 1975*), which show remarkably preserved soft parts. We undertake a detailed morphological analysis of this new material to assess the hypotheses of the possible affinity and lifestyle of this bivalved taxon, and its implications for understanding the diversity and evolution of early euarthropods

## Results

### Systematic palaeontology

Phylum: Euarthropoda Lankester, 1904

### Genus (Monotypic)

*Chuandianella* Hou and Bergström, 1991

### Generic and species diagnosis (amended after Hou and Bergström)

Bivalved euarthropod bearing a uniramous antennule consisting of at least ten podomeres; a short, uniramous second appendage with at least six podomeres; ten other, homonomous appendages, each comprising a short paddle-shaped exopod and a feather-like endopod bearing at least 27 podomeres each with a long blade-like endite bearing a terminal seta; and an abdomen comprising four apodous segments, plus a tailpiece with two elongate, flap-like caudal structures.

### Type species

*Mononotella ovata* Li, 1975

Note: The author’s name “Li” is spelled “Lee” in some publications.

### Type horizon and locality

Chiungchussu Formation, *Eoredlichia*-*Wutingaspis* trilobite biozone, Cambrian Series 2, Stage 3. Chiungchussu, Kunming, Yunnan Province, China.

### Species

*Chuandianella ovata* (Li, 1975).

### Holotype

Repository given (*Li, 1975)* as the Institute of Southwestern Geosciences, Chengdu, China (now known as The Chengdu Centre of the Geological Survey of China). Collection number YN6303, specimen number YO10; designated and figured Li 1975, pl. 3, fig. 16.

### Key synonymy

*Mononotella ovata* Lee, 1975, sp. nov., p. 65, pl. 3, figs 16, 17*; Mononotella viviosa* Lee, 1975, sp. nov., p. 65, pl. 3, fig. 18; ?*Mononotella marginia* Lee, 1975, sp. nov., pl. 3, figs 19, 20; *Chuandianella ovata* (Li, 1975), Hou and Bergström, 1997, p. 41, fig. 37 (q.v. for earlier synonymy); *Chuandianella ovata* (Li, 1975), Liu and Shu, 2008, p. 358, text-figs 1–3; *Chuandianella ovata* (Lee, 1975), Hou *et al*., 2017, p. 238, figs 20.61, 20.62; *Chuandianella ovata*, Ou *et al*., 2020, figs 1A,C,E,G,H,J, 2, 3.

**Figure 1.**
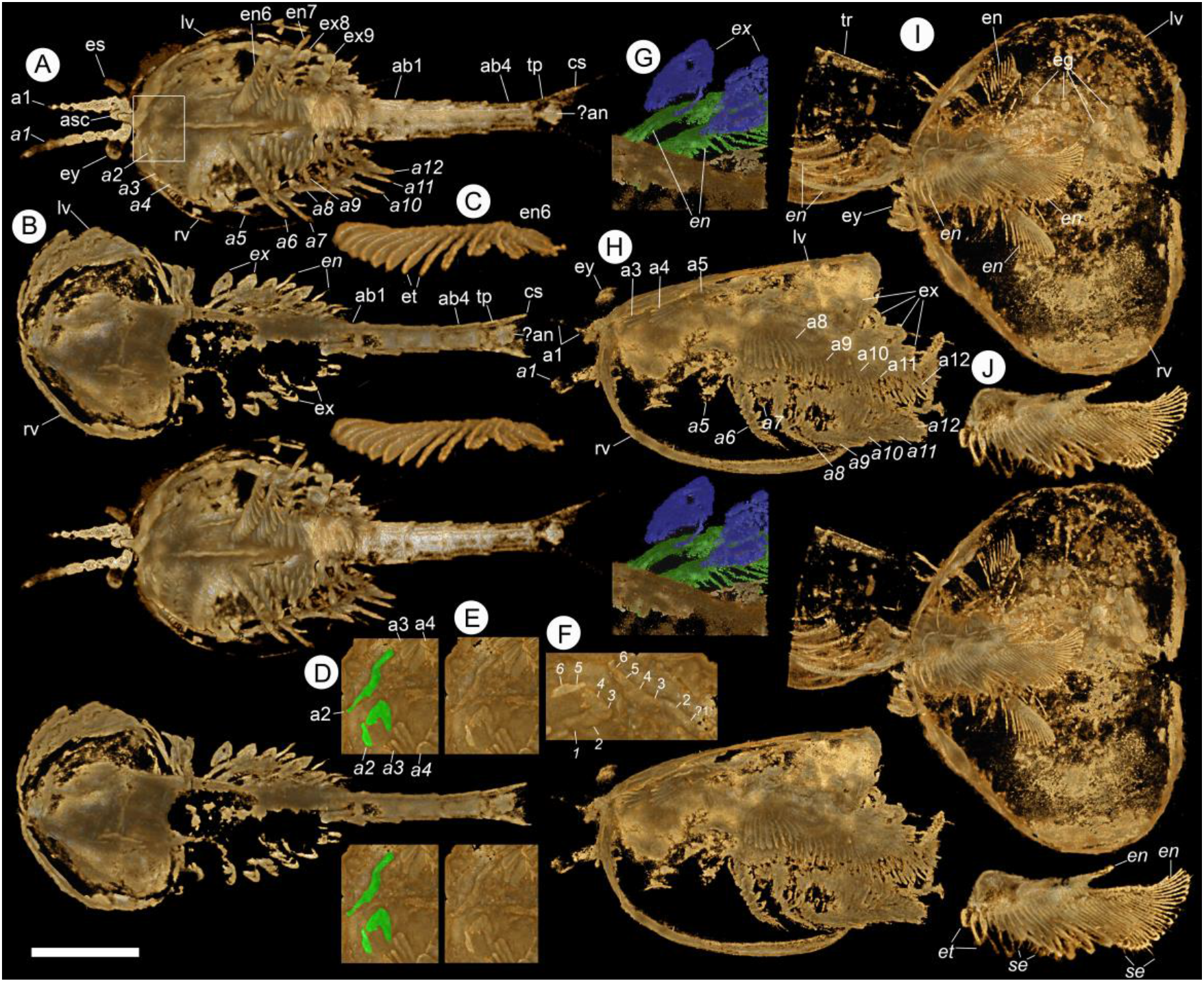
Micro-CT images of *Chuandianella ovata*. (A,C-F) YKLP 16218. (A) Ventral view. Scale bar = 5.0 mm. (C) Endopod of 6^th^ appendage showing endites. Scale bar = 1.6 mm. (D) Ventral view (white rectangle in A), showing details of second appendage (a2, green). Scale bar = 2.0 mm. (E) Same as D but with a2 not coloured. Scale bar = 2.0 mm. (F) Close-up view of 2^nd^ appendages (rotated by 90 degrees with anterior to upper position), with podomeres numbered. Scale bar = 1.4 mm. (B,G) YKLP 16216. (B) Taphonomically deformed specimen, showing ventral view of the carapace and dorsal view of the soft parts (cf. ***Supplementary Figure 1***). Scale bar = 3.8 mm. (G) Dorsal view of part of right side of the body, showing exopods (blue) and endopods (green). Scale bar = 1.4 mm. (H) YKLP 16238, left lateral view, showing endopods and exopods of trunk appendages. Scale bar = 4.9 mm. (I,J) YKLP 16239. (I) Ventral view, showing *circa* 20 eggs within the left valve. Scale bar = 4.3 mm. (J) Endopods of 3^rd^ and 5^th^ (?) appendages showing long blade-like endites each with a terminal seta. Proximally in this image two endopods overlap, giving the false impression of setae along the lateral margins of the endites. Scale bar = 3.4 mm. All panels are stereo-pairs. Abbreviations: a1, antennule; a2, second appendage; a3-a12, biramous appendages; ab1-4, abdominal segments 1 to 4; an, anus; asc, anterior sclerite; cs, caudal structure; eg, egg; en, endopod; es, eye stalk; et, endite; ex, exopod; ey, stalked eyes; lv, left valve; rv, right valve; se, seta; tp, tailpiece; tr, trunk. Italics indicate a right-side appendage.

**Figure 2.**
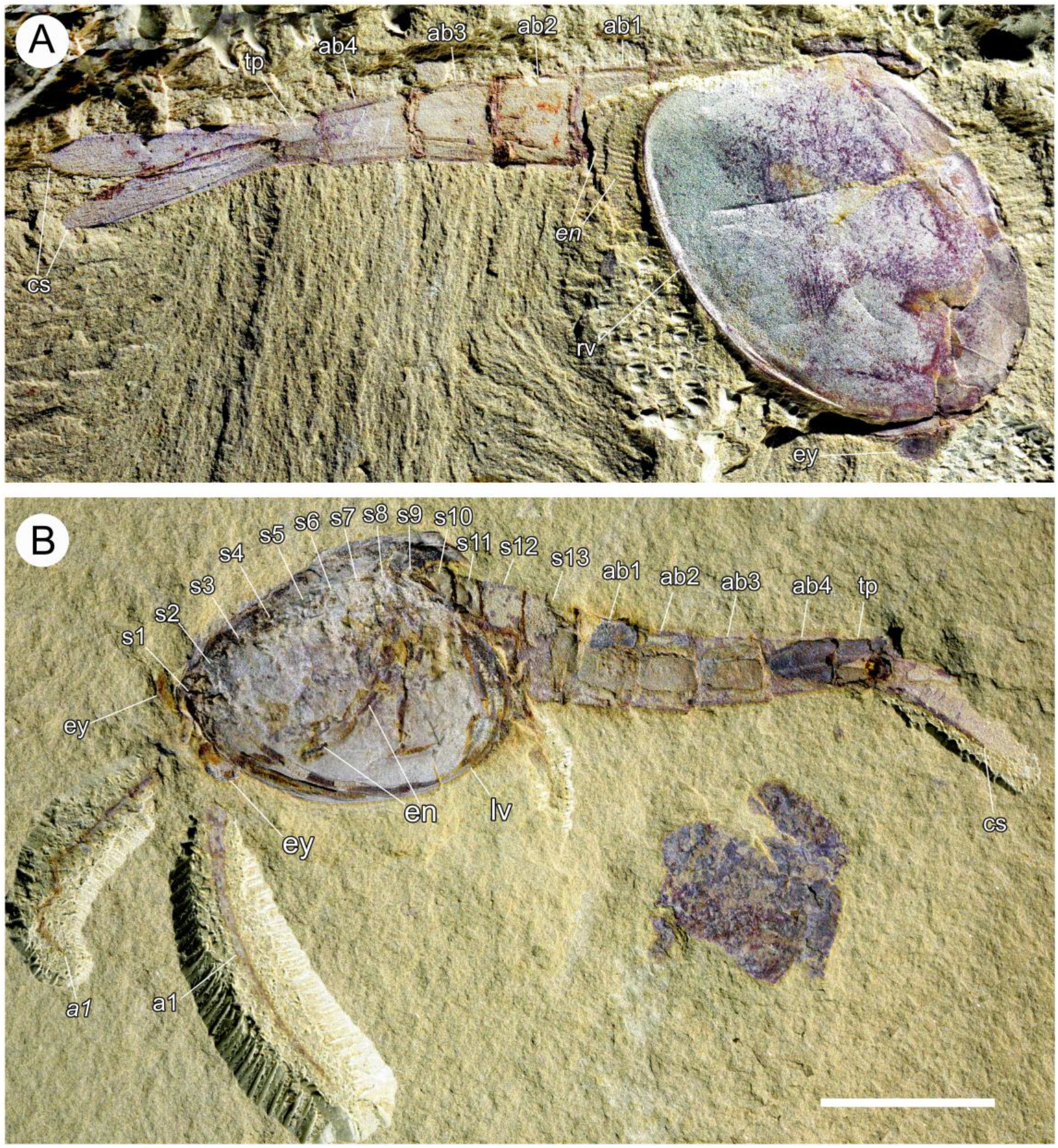
Photographs of *Chuandianella ovata*, showing overall morphology and segmentation of the body. (A) RCCBYU 10272, right lateral view. Scale bar = 3.4 mm. (B) YKLP 13967a, left lateral view. Scale bar = 5.0 mm. Abbreviations additional to fig. 1: s1-s12, head and thoracic segments. s1 is the eye-bearing segment/anterior sclerite; the position of segments s1 and s2 is difficult to infer. Italics indicate a right-side appendage.

**Figure 3.**
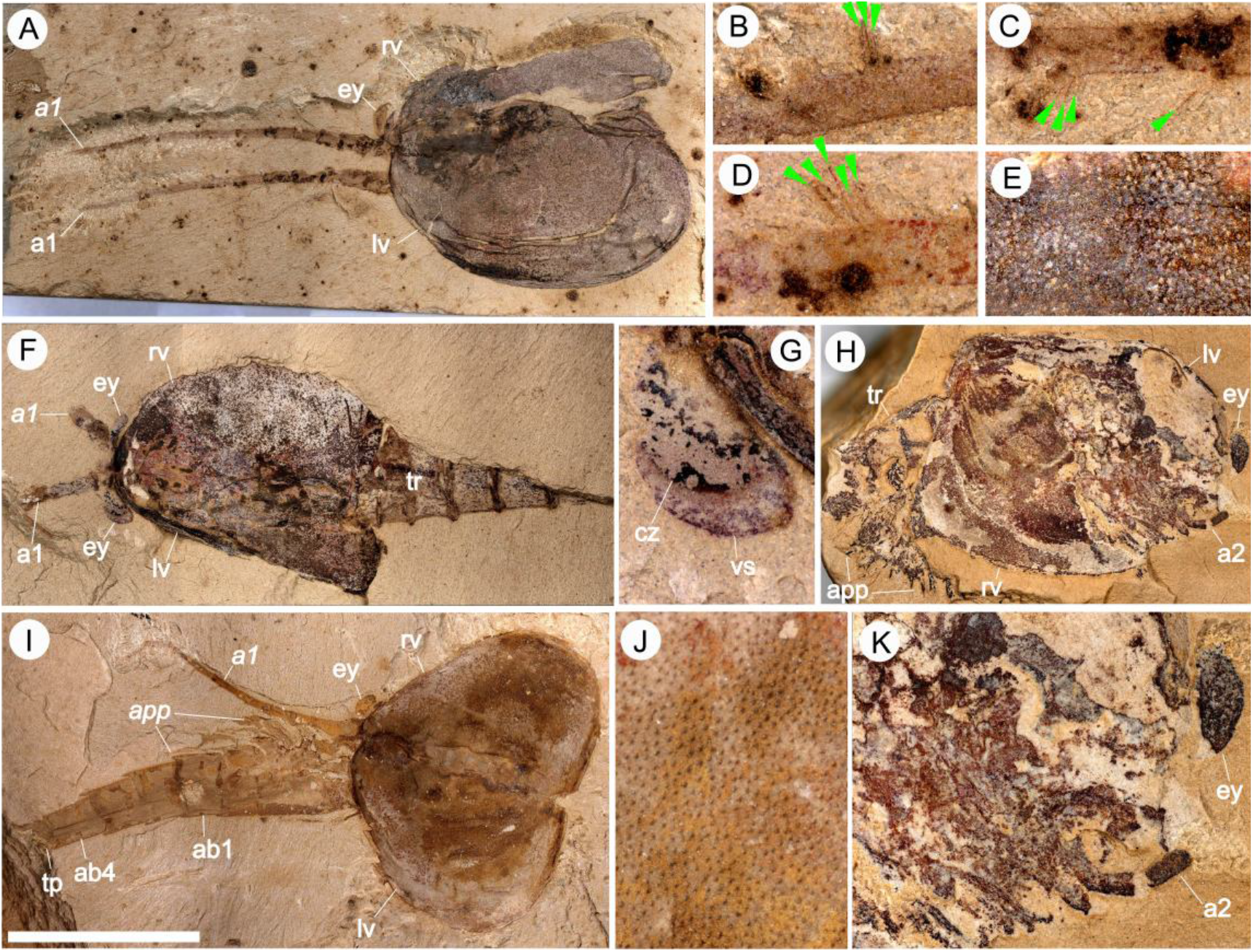
Photographs of *Chuandianella ovata*, showing morphological details. (A-E) YKLP 16217a (cf. ***Supplementary Figure 3***). (A) Dorsal view. Scale bar = 8.3 mm. (B) Setae on inner surface of 7^th^ podomere of left antenna. Scale bar = 0.1 mm. (C) Setae on inner surface of 5^th^ podomere of right antenna. Scale bar = 1.6 mm. (D) Setae on inner surface of 5^th^ podomere of left antenna. Scale bar = 1.3 mm. (E) Pits on carapace surface. Scale bar = 1.0 mm. (F,G) YKLP 16238a, specimen depicted in ***Figure 1f***. (F) dorsal view. Scale bar = 8.4 mm. (G) Visual surface of the left eye. Scale bar = 1.6 mm. (H,K) YKLP 16259, a laterally compressed specimen with anterior part of right valve missing, exposing the anterior appendages. Posterior part of trunk is also missing. (H) Overview. Scale bar = 6.0 mm. (K) Detailed view of anterior part, showing position and morphology of a2. Scale bar = 2.4 mm. (I,J) YKLP 16239, specimen depicted in ***Figure 1g,h***. (I) Dorsal view with trunk reflexed so that it appears to emerge from anterior end of carapace. Scale bar = 8.8 mm. (J) Pits on carapace surface. Scale bar = 0.9 mm. Abbreviations additional to figs 1, 2: app, appendage; vs, visual surface. Italics indicate a right-side appendage.

### Species diagnosis

As for the genus.

### Description

The carapace is up to 1.45 cm long (*Liu and Shu, 2008*), ‘bivalved’ along a median fold, but lacking an articulating hinge. Valves are strongly postplete in lateral outline and lack lobation; they have a narrow incurved free margin (*Figures 1h, 2a,b, 4a*). In some individuals *(Figure 3e, j; Supplementary Figure 7a)* the external carapace surface is finely pitted. The body is up to 3 cm long (*Ou et al., 2020*) and consists of 18 segments (*Figures 2b, 4*). It is attached to the carapace dorsally by at least the first four (possibly five) segments (*Figure 1h*). Pedunculate stalked eyes originate from the first (ocular; presumed protocerebral) segment and protrude beyond the anterior margin of the carapace (e.g., *Figure 3f,g; Supplementary Figures 4, 6*). The eye has a dark-coloured central zone and a light-coloured outer zone, with a well-defined visual surface (*Figure 3g*; *Supplementary Figure 6b*).

**Figure 4.**
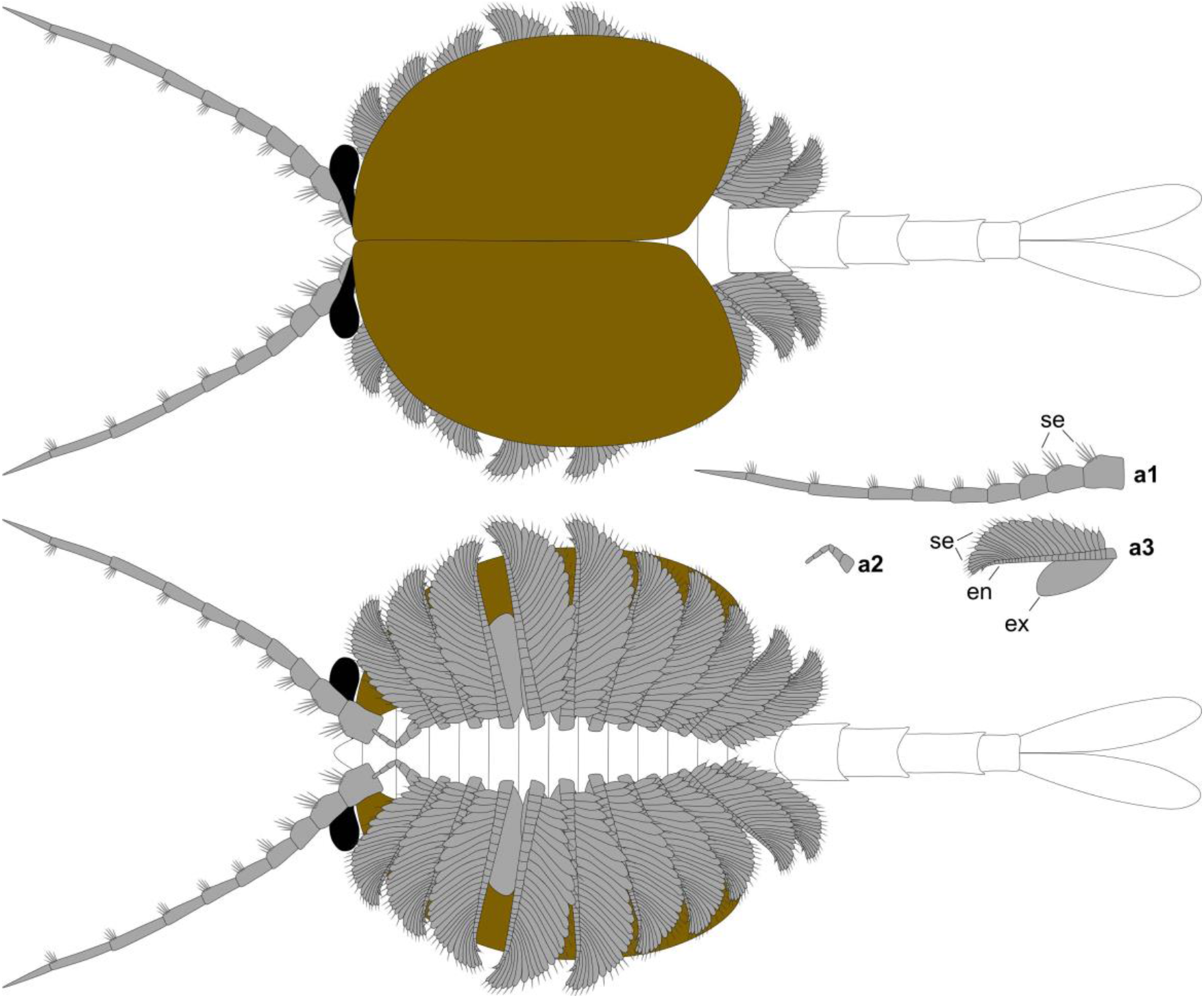
Reconstruction of *Chuandianella ovata*. Upper: dorsal view. Lower: ventral view. Right middle: isolated appendages a1-a3. Not to scale. Abbreviations as in figs 1-3.

The antennule is uniramous, narrow and about 30% longer than the carapace; it consists of at least 10 podomeres and gradually tapers distally (*Figures 1a, 2b, 3a,i*; *Supplementary Figures 5a, 6a*). The proximal podomere is stouter than the rest and the more distal podomeres are longer; each podomere bears up to five short stiff setae on the adaxial surface (*Figure 3b-d; Supplementary Figure 5*). The second appendage is uniramous, consisting of at least six, gradually tapering podomeres; it is presumed to represent an endopod (*Figures 1d,e,f, 3k, 4*; *Supplementary 8b,c*). Its terminal podomere is elongate, rod-like and apparently lacks a terminal claw (*Figures 1f, 3k*); it is uncertain if it has setae or not (*Supplementary Figure 8c*). Posterior to the second appendage there are ten homonomous appendages. Each consists of a short paddle-shaped exopod (*Figures 1b,g*; *Supplementary Figure 1a,b, 2a,b, 7d*) and a long endopod that is more robust proximally and gradually tapers distally (*Figure 1j*); evidence of the basipod is not apparent. The endopod bears at least 27 podomeres, each with a long blade-like endite bearing a short terminal seta (*Figure 1c,j, 4*; *Supplementary Figures 2c, 3b, 5b*), giving an overall feather-like morphology. The endites in some specimens are preserved perpendicular to the axis of the endopod and parallel to each other (e.g., *Figure 1c,h*), but in other specimens they overlap each other (e.g., *Figure 1j*) indicating flexible movement and/or taphonomic displacement. The posterior part of the trunk (= abdomen *sensu Vannier et al., 2018* and *Zhai et al., 2019b*), which consists of a tubular section of four sclerites and a tailpiece bearing two long, blade-like structures, is apodous (*Figures 1a,b, 2a,b*). The carapace covers about the first nine segments of the body; the more posterior segments protrude posteriorly from the carapace via a gape (*Figure 2a,b*). Two specimens bear tiny sub-circular/ovoid objects, mostly 400–600 μm across and loosely scattered within the left valve (*Figures 1j*) or roughly arranged in multiple rows (*Supplementary Figure 9a*); they are interpreted as eggs. In the better-preserved specimen YKLP 16258 (*Supplementary Figure 9*) at least 48 eggs are associated with a single valve.

**Figure 5.**
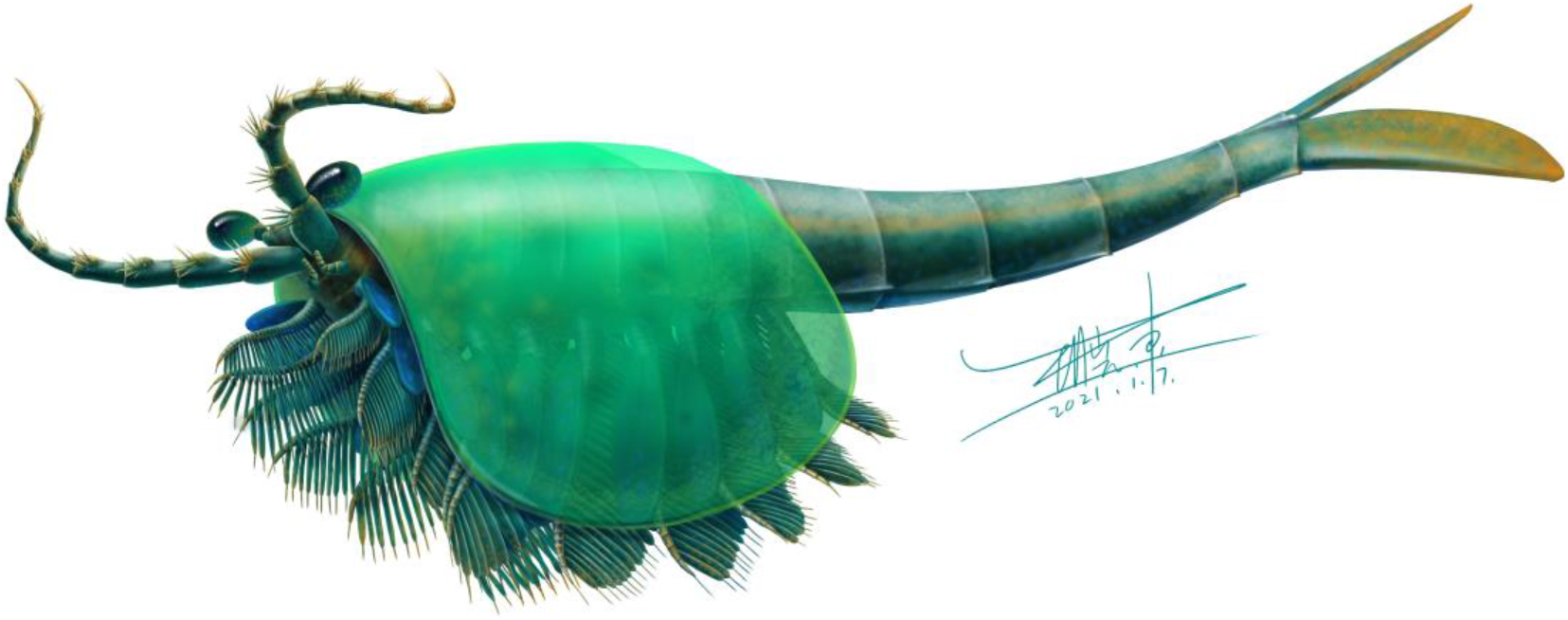
Reconstruction of *Chuandianella ovata in vivo*.

## Discussion

### *Chuandianella ovata* is not a waptiid

Based on carapace morphology *C. ovata* was originally (*Li, 1975*) assigned to the Cambrian bradoriid *Mononotella* (for which, see *Siveter and Williams, 1997*) and was subsequently designated (*Hou and Bergström, 1991*) as the type species of *Chuandianella*. *Chen et al. (1996)* and Chen *(2004)* opined that *C. ovata* is a waptiid, related to the Burgess Shale *Waptia fieldensis* Walcott, 1912. Hou and Bergström*, (1997)* tentatively included *Chuandianella* in the Family Waptiidae Walcott, 1912, though noted it had ‘not yet been studied in detail’. Both taxa have carapaces with a postplete outline lacking lobes and nodes, a dorsal median fold without an articulating hinge, a trunk with four apodous segments and a tailpiece with two caudal structures (*Chen et al., 1996; Hou and Bergström, 1997*; *Chen, 2004*). With some exceptions (*Liu and Shu, 2004, 2008*; *Hou et al. 2017)* a possible affinity with waptiids has persisted, as in the study of the bivalved *Pauloterminus spinodorsalis* from the Cambrian of Greenland (*Taylor, 2002*) and in *Vannier et al. (2018*) who considered *Chuandianella* to be a mandibulate euarthropod.

*C. ovata* differs in morphology from both *P. spinodorsalis* and *W. fieldensis*. The biramous trunk appendages of *P. spinodorsalis*, each with a short feather-like endopod and a long, paddle-shaped exopod, distinguish it from *C. ovata,* whilst *P. spinodorsalis* also possesses a longer carapace (9.1 - 46.3 mm; *Taylor, 2002*) compared with that of *C. ovata* (5.15 - 13.09 mm; see Table 1)*. Waptia fieldensis* is interpreted as possessing a specialised mandible and maxillula, whilst the four post-maxillular appendages are also specialized with 5-segmented endopods (*Vannier et al., 2018*). These important and diagnostic features, together with the longer carapace of *W. fieldensis* (in adults,10.99 - 24.54 mm; *Vannier et al., 2018*), distinguish it from *C. ovata*.

The morphology of *Chuandianella* does not support its assignment to crown-group Mandibulata, not least because of the absence of head segments bearing mandibles and maxillulae (see *Scholtz and Edgecombe 2006*). Here we interpret *Chuandianella* as an ‘upper’ stem-group euarthropod (*sensu Ortega-Hernández, 2016*) based on its possession of a deutocerebral first appendage pair, a multi-segmented head region with, in this case, two pairs of differentiated post-ocular limbs, complete arthropodization, including post-oral biramous limbs. We cannot determine the presence of a posterior facing mouth beneath a hypostome/labrum complex.

### The feather-like endopods of *C. ovata* are unique

Previous studies of *C. ovata* (e.g., *Liu and Shu, 2008*) failed to identify the paddle-shaped exopods of its trunk limbs, and the feather-like endopods (*Figures 1c,j*; *2a*) were mistakenly interpreted as exopods. The extremely long blade-like endopodal endites of *C. ovata* are the only ones of their kind known in a Cambrian bivalved euarthropod and further demonstrate that early, ‘upper’ stem-group euarthropods were experimenting with a wide array of different limb arrangements and morphologies (*Zhai et al., 2019a*). Possible comparable structures occur in *W. fieldensis*: six pairs of annulate post-cephalothoracic seemingly single-branched appendages fringed with long lamellae (*Vannier et al., 2018)* were interpreted as possible endopods or, more likely, basipods of a unique morphology within Euarthropoda (*Vannier et al., 2018)*. The morphology of the endopods in *C. ovata* also bear comparison with those of the bradoriid *Kunmingella douvillei* (Mansuy, 1912), though in the latter the endopodal endites are less numerous *(see Zhai et al., 2019a)*, are cylindrical rather than blade-like, and are significantly shorter. Without a full understanding of their 3-D morphology, the endopods of *Kunmingella* were initially misinterpreted as exopods (*Hou et al., 1996*, *fig. 5*).

### The bivalved carapace is an unreliable indicator of affinity

Previous studies have clearly demonstrated that the taxonomic assignment of early Palaeozoic ostracod crustaceans may be flawed if based on the morphology of their bivalved carapace alone (*Siveter et al., 2012, 2018*). Similarly, analysis of three bivalved arthropods referred to the Bradoriida has demonstrated that their carapace morphology when considered alone is an unreliable basis for classification: the carapace houses markedly different soft-bodies that include both stem-euarthropods and mandibulate-like euarthropods (*Zhai et al., 2019a*). The data presented here on the soft-part anatomy of *C. ovata* provides further evidence that carapace morphology alone is a poor indicator of the affinity of bivalved euarthropods and that the diversity of morphologies seen in bivalved fossil euarthropods is greater than previously appreciated.

### Mode of life of *C. ovata*

The morphology of the biramous appendages of *C. ovata* is not compatible with ambulatory activity on the seabed. The feather-like endopods and well-developed tail fan of *C. ovata* may have aided swimming/propulsion and manoeuvrability (*Figures 1c,j, 4, 5*; see also *Liu and Shu, 2008).* A possible nektonic lifestyle is also supported by its occurrence: *C. ovata* is relatively common and is widespread in the Cambrian of southwest China (*Hou et al., 2017*). It is known from similar Cambrian stratigraphical levels (Series 2, Stage 3) in Sichuan, Guizhou and southern Shaanxi provinces (although only material from Yunnan has yielded soft-part anatomy). *C. ovata* may have used its long feather-like endopodal endites for filter-feeding, capturing small-sized organic material. Its long setate antennules presumably had a sensory function, perhaps to detect predators, food or monitor environmental conditions. The diminutive second appendage may have functioned like the main ramus of the first maxillula of living crustaceans such as ostracods (*Meisch*, 2000) to support food manipulation other than mastication. The stalked eyes are well developed, protrude beyond the carapace and their preservation in various orientations suggests that they were mobile to provide multi-directional vision (*Figures 1a, 2a,b, 3a,f,i; see also Supplementary Figures 4,6a*). The radius of curvature of the eye is greater laterally than frontally, suggesting better resolution of the lateral field (*Strausfeld*, 2015). That *C. ovata* occurs in supposed coprolites in the Chengjiang biota (*Chen and Zhou, 1997; Vannier and Chen, 2005*) indicates that it was a prey or carrion item.

The bivalved carapace of *C. ovata* apparently functioned not only for protection of soft parts but also apparently as a surface for the attachment of its eggs. *Ou et al. (2020)* reported egg-bearing specimens of *C. ovata* and compared possible reproductive modes of *C. ovata* and *W. fieldensis* (the latter reported by *Caron and Vannier, 2016*) based on the size, number and morphology of eggs. They determined that the eggs of *C. ovata* were smaller (0.5 mm versus 2.0 mm in diameter) than those of *W. fieldensis* but each individual animal carried significantly more eggs (≤100 versus ≤26 per clutch) than *W. fieldensis*, implying different reproductive strategies (*Ou et al., 2020*). Our observations on our egg-bearing specimens of *C. ovata* (*Figure 1i; Supplementary Figure 9*) generally confirm the size, number and position of eggs as indicated by *Ou et al. (2020)*. Since *C. ovata* is morphologically distinct from *W. fieldensis* differences in brooding strategies between these taxa are not surprising. Sexual dimorphism has been suggested for *C. ovata*, by which the valves of supposed males are larger, with a greater height to length ratio, and have a pitted rather than smooth surface (*Liu and Shu, 2008)*. As only one of the two egg-bearing specimens in our material has pitted valves (*Figure 3j*) ornament should not be regarded as a possible dimorphic character. We have been unable to replicate the observation (*Liu and Shu, 2008*) that female and male reproductive systems are preserved in some specimens.

### Conclusions

Micro-CT scanning of the stem group euarthropod *Chuandianella ovata* from the Cambrian Chengjiang Lagerstätte reveals unprecedented details of its soft-part anatomy. Notably, *C. ovata* possessed differentiated first and second appendages, and a further ten homonomous appendages each with a short paddle-shaped exopod and a feather-like endopod of at least 27 podomeres. This morphology clearly differentiates *Chuandianella* from the Cambrian mandibulate euarthropod *Waptia*, to which it has been consistently compared.

The feather-like endopods of *C. ovata* attest to the wide diversity of limb arrangements and morphologies developed by early Cambrian, ‘upper’ stem-group euarthropods. Together with the well-developed tail fan of *C. ovata,* these may have facilitated a nektonic lifestyle, a notion that is also supported by the widespread occurrence of *C. ovata* in the Cambrian of southwest China. Its well-developed stalked eyes would have provided multi-directional vision for various uses including detection of predators. That *C. ovata* occurs in supposed coprolites in the Chengjiang biota also indicates that it was a prey or carrion item.

## Material and methods

New specimens of *C. ovata* were collected from the Yu’anshan Member, Chiungchussu Formation, *Eoredlichia*-*Wutingaspis* trilobite biozone, Cambrian Series 2, Stage 3, Yunnan Province (see *Hou et al., 2017*), at Mafang, Ercaicun and Jianshan in Haikou, Kunming (*Supplementary Table 1*). Fourteen specimens which revealed appendage morphology in high fidelity were selected for detailed study. Specimens are mainly housed in the Yunnan Key Laboratory for Palaeobiology (YKLP), Yunnan University, Kunming, or in the Yunnan Geological Survey (Hz-f-4-777, He-f-6-4-294).

Fossil structures exposed on the surface of the rock slabs were imaged with a Nikon D3X camera with an Af-S VR105 macro lens (*Figure 3*) and a Keyence VHX6000 stereo-microscope (photographs in all other figures). Fossil structures hidden within the slabs (*Figures 1, 2*) were revealed using a Zeiss Xradia 520 Versa X-ray Microscope. Scanning pixel size ranged from 3.4 to 26.8 μm, depending on the size of the scanned region and the slab. The digital data from each specimen, in the form of a series of one to a few thousand TIFF images representing cross-sections through different parts of the slab, were processed with Drishti software (Versions 2.4) to generate 3-D models of the fossils.

## Acknowledgements

This study was supported by NSFC grants 41861134032 and 41902011, the Key Research Program of the Institute of Geology & Geophysics, Chinese Academy of Sciences (IGGCAS-201905), and Yunnan Provincial Research Grant YNWR-QNBJ-2019-295. M.W. thanks the Leverhulme Trust for a Research Fellowship (RF-2018-275\4). We thank Mr. Xiaodong Wang for making the artistic reconstruction of *Chuandianella* used in Figure 5. The Jianshan Subsidiary of the Yunnan Phosphate Chemical Group Co. Ltd. provided invaluable help facilitating field work. The Yunnan Geological Survey granted access to two of the specimens.

## Additional files

### Supplementary files

Supplementary Table 1. Dimensions of specimens of *Chuandianella ovata* investigated in this study.

Supplementary Figure 1. *Chuandianella ovata*, YKLP 16216. (A) Microscope image of specimen on rock slab. Scale bar = 0.8 mm. (B) Micro-CT stereopair image. Scale bar = 2.0 mm. Abbreviations as for Figures 1–3. Italics indicate a right-side appendage. Note: The soft parts of this specimen are taphonomically dislocated and are flipped vertically, so that this figure shows the dorsal views of the carapace and the ventral views of the trunk and appendages, with the left appendages associated with the right valve while the right appendages are associated with the left valve (cf. *Figure 1b,g*).

Supplementary Figure 2. *Chuandianella ovata*, YKLP 16215a. (A) Microscope image of specimen on rock slab, dorsal view. Note that the posterior part of the trunk is missing. Scale bar = 3.3 mm. (B,C) Micro-CT images of anterior part, stereo-pairs (anterior end to the left). Scale bar = 3.1 mm. (B) Dorsal view. (C) Ventral view. Abbreviations as for Figures 1–3. Italics indicate a right-side appendage.

Supplementary Figure 3. *Chuandianella ovata*, YKLP 16217 (for microscope image of this specimen see *Figure 3a-e*), stereo-pairs of micro-CT images. Scale bar = 5.0 mm. (A) Dorsal view of anterior part. (B) Ventral view of anterior part. Abbreviation additional to Figures 1–3: cz, central zone of the eye with dark coloration. Italics indicate a right-side appendage.

Supplementary Figure 4. *Chuandianella ovata*, YKLP 16218, microscope images (for micro-CT images of this specimen see *Figure 1a, c*). Scale bar = 1.0 mm. (A) YKLP 16218a, dorsal view of anterior part. (B) YKLP 16218b, ventral view of anterior part; posterior part is buried in sediment. Abbreviations as for Figures 1–3. Italics indicate a right-side appendage.

Supplementary Figure 5. *Chuandianella ovata*, He-f-6-4-294. (A) Microscope image, oblique-right view; the posterior part of the trunk is missing. Scale bar = 1.0 mm. (B) Stereo-pair of micro-CT image, details of anterior part of specimen, oblique-left view (viewed from underside of slab). Scale bar = 1.2 mm. (C) Details of setae on left a1 (white rectangle in A). Scale bar = 0.5 mm. Abbreviations as for Figures 1–3. Italics indicate a right-side appendage.

Supplementary Figure 6. *Chuandianella ovata*, Hz-f-4-777. (A) Microscope image, dorsal view; the left and posterior parts of the specimen are missing. Scale bar = 1.8 mm. (B) Details of left eye. Scale bar = 0.7 mm. (C) Stereo-pair of micro-CT image, ventral view of anterior part. Scale bar = 1.0 mm. Abbreviations as for Figures 1–3. Italics indicate a right-side appendage.

Supplementary Figure 7. *Chuandianella ovata*, YKLP 16256. (A) Microscope image, showing pits on left valve. Scale bar = 0.4 mm. (B) Microscope image, dorsal view. Scale bar = 0.9 mm. (C,D) Stereo-pairs of micro-CT images. The posterior part of the body is reflexed so that it emerges from the anterior end of the carapace. Scale bar = 2.0 mm. (C) Dorsal view. (D) Ventral view. Abbreviations as for Figures 1–3. Italics indicate a right-side appendage.

Supplementary Figure 8. *Chuandianella ovata*, YKLP 16257. (A) Microscope image, dorsal view. Scale bar = 2.3 mm. (B,C) Stereo-pairs of micro-CT images. (B) Dorsal view. Scale bar = 5.0 mm. (C) Ventral view of the anterior part (white rectangle in B), showing a2. Scale bar = 1.5 mm. Abbreviation additional to Figures 1–3: uf, unidentified fossil. Italics indicate a right-side appendage.

Supplementary Figure 9. *Chuandianella ovata*, YKLP 16258, an egg-bearing specimen, microscope images. (A) Overview, oblique-right view. Scale bar = 2.0 mm. (B) Details of eggs (white rectangle in A). Scale bar = 1.4 mm. Abbreviations as for Figures 1–3. Italics indicate a right-side appendage.

## Data availability

Computed tomography data will be available on Dryad upon acceptance by the journal.

